# A transition zone enriched *WIF1*⁺ basal cell subtype is associated with benign prostatic hyperplasia

**DOI:** 10.64898/2025.12.08.692826

**Authors:** Rulin Wang, Qizhi Zheng, Mindy K Graham, Ajay Vaghasia, Jianyong Liu, Jordan Gregg, Tracy Jones, Anuj Gupta, Nicole Castagna, Yan Zhang, Kornel Schuebel, Jennifer Meyers, Alyza Skaist, Dixie Hoyle, Yuhan Yang, William G Nelson, Angelo M De Marzo, Srinivasan Yegnasubramanian

## Abstract

The cellular composition and disease susceptibilities of the distinct zones of the human prostate remain incompletely understood. Through extensive single-cell RNA sequencing (scRNA-seq) of benign regions from prostatectomy specimens, we identified a basal cell population expressing *WIF1, VCAN*, and *NRG1*, among other genes, that was significantly enriched in the transition zone (TZ). Benign prostatic hyperplasia (BPH) is a common condition that causes widespread morbidity and is nearly exclusively localized to the TZ. Analysis of previously published scRNA-seq datasets further confirmed that *WIF1*+ basal cells were significantly enriched in BPH compared to normal prostate. Pathway and cell-cell communication analyses revealed that this basal subtype is associated with programs related to cell proliferation, epithelial-mesenchymal transition (EMT), angiogenesis, and hormone response. Together, the molecular signature, zonal distribution, and pathway enrichment suggest that TZ-enriched *WIF1*+ basal cells may contribute to BPH pathogenesis by promoting epithelial and stromal remodeling.

## INTRODUCTION

The human prostate is an androgen-regulated organ anatomically divided into distinct zones- peripheral zone (PZ), central zone (CZ), transition zone (TZ), as well as the anterior fibromuscular stroma, which lacks epithelium. Each of the epithelial rich zones (PZ, CZ and TZ) harbor at least some unique histological features, cellular composition, and disease susceptibilities. Notably, benign prostatic hyperplasia (BPH) arises almost exclusively in the TZ, whereas prostate cancer develops predominantly in the PZ [1]. This striking zonal selectivity underscores the need to understand spatial heterogeneity in prostate biology and pathology.

Recent advances in single-cell RNA sequencing (scRNA-seq) have enabled dissection of the prostate’s complex architecture at unprecedented resolution [2–5]. These studies have uncovered substantial heterogeneity within both epithelial and stromal lineages, revealing zone-specific cell states, lineage trajectories, and molecular signaling pathways. For example, Henry et al. identified two novel epithelial populations, termed club and hillock cells, which were more abundant in the TZ than in the PZ [2]. Yan et al. further demonstrated age-associated changes, reporting a *TFF3*+ luminal subtype with elevated *MYC* and *E2F* activity enriched in the PZ, while the TZ exhibited stronger transcriptional signatures related to immunity and stemness [3]. Stromal heterogeneity has also been noted, with Joseph et al. and Yan et al. both describing proximal-distal fibroblast density differences [3,4]. Despite these insights, basal cell heterogeneity has received limited attention and remains poorly understood, likely due to small sample sizes [5]. One recent study described two lineages of *KRT5*+ basal cells (*KRT16*+*KRT17*+ and *KRT16*^-^*KRT17*^-^), which were further subdivided into seven transcriptionally distinct subtypes [6]. The other study described multiple basal subpopulations, including *KRT15*+/c1 and *DST*+/c6 clusters representing a classical basal phenotype, a rare *RARRES2*+/c17 basal subtype, and *S100A2*+/c2 and *PLCG2*+/c3 subtypes that were enriched for basal and hillock cell signatures [7]. Most prior studies focused solely on normal prostate tissue, limiting their ability to connect cell states with disease contexts such as BPH.

BPH is characterized by non-malignant proliferation of epithelial and stromal cells, leading to prostatic enlargement that can lead to urethral obstruction, causing lower urinary tract symptoms (LUTS) that substantially reduce quality of life. Although the pathogenesis of BPH remains incompletely understood, dysregulated androgen signaling [8], chronic inflammation [9], and stromal-epithelial interactions [10] are considered as key contributors. Additional studies have implicated cellular senescence, extracellular matrix remodeling, and hormonal imbalances in disease progression [11,12].

In this study, we analyzed peri-cancerous prostate tissues from ten patients spanning all three anatomical zones and identified a distinct basal cell population-*WIF1*+ basal cells. This subtype was significantly enriched in the TZ and in BPH tissues, pointing to a zonally restricted population potentially involved in BPH pathogenesis. These findings provide a potential novel cellular basis for the clinical observation that prostate cancer arises predominantly in the peripheral zone, whereas BPH develops almost exclusively in the transition zone.

## RESULTS

### Identification of Major Cell Types in Human Prostate

We performed single-cell RNA sequencing (scRNA-seq) on prostate tissues from ten individuals who underwent radical prostatectomy for treatment of localized prostate cancer. From each subject, benign regions were sampled from the peripheral (PZ), central (CZ), and transition (TZ) zones (Fig. 1A, 1B). After excluding low-quality samples due to technical failures (e.g., wetting failure), we retained seven PZ, nine CZ, and nine TZ samples, yielding a total of 129,887 high-quality single cells across 25 samples. Data from the peripheral zone were previously reported in the context of assessing tumor specific alterations [13], and were re-analyzed with the data from the other two zones in this study.

**Figure 1.**
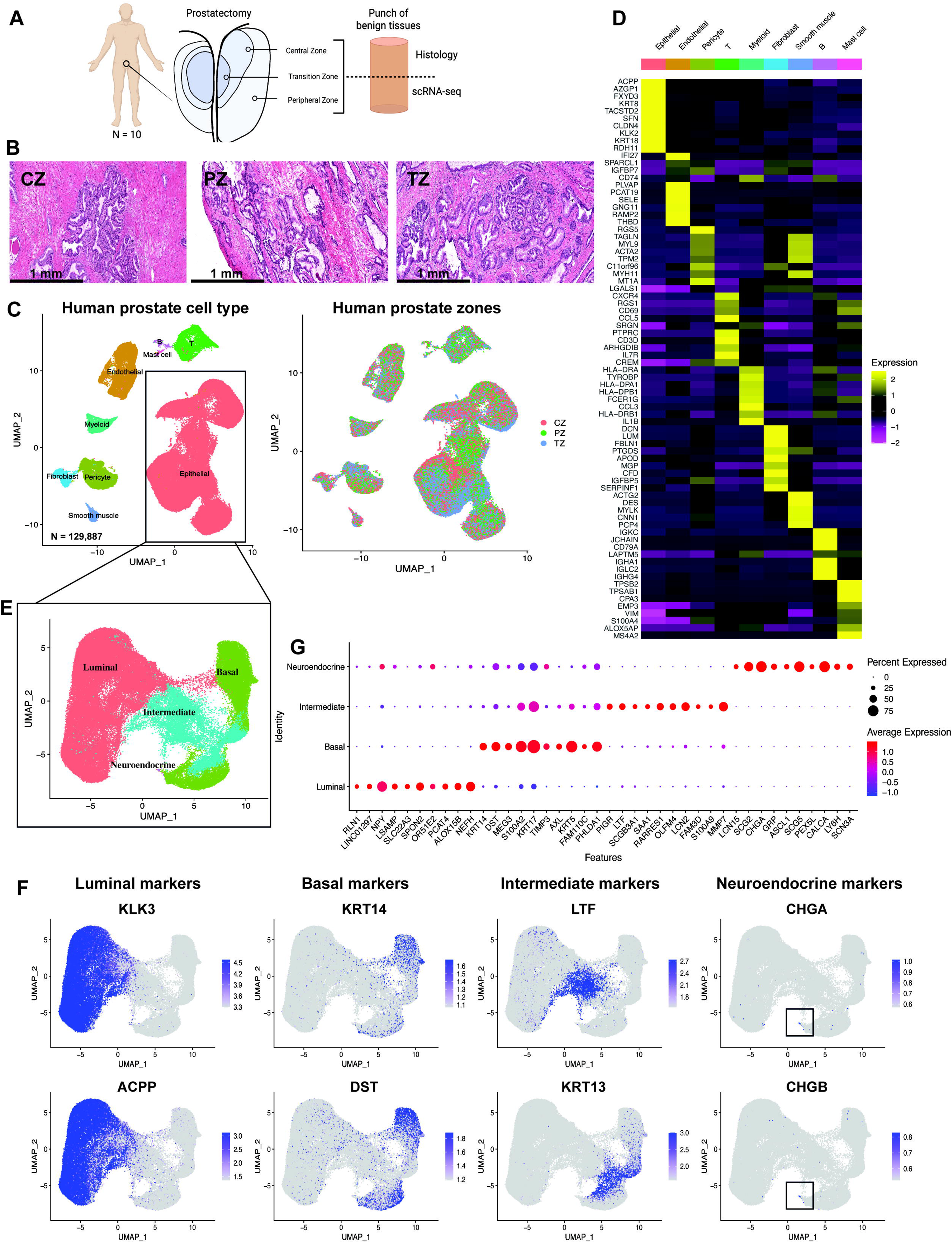
Single cell sequencing and histopathology of zone-specific tissues collected from cancer-free regions of prostatectomies. A. For each prostatectomy (*n* = 10), tissue punches were collected from each zone of the prostate (three punches per subject corresponding to each of the three zones). For each tissue punch, part of the tissue was processed for scRNA-seq from freshly dissociated cells, and the remaining half was frozen and sectioned for histology, and in situ hybridization. B. Representative examples of hematoxylin and eosin (H&E) staining of fresh frozen central zone (CZ), peripheral zone (PZ), and transition zone (TZ) tissue punches. Scale bar indicates 1 mm. C. Dimensionality reduction (uniform manifold approximation and projection, UMAP) and clustering analysis of scRNA-seq of >129,000 cells from CZ, PZ, and TZ tissue showed cell clustering by known cell types across the three zones. D. Heatmap of the top 10 differentially expressed genes of each cell type. Sorted by log2FC for each cell type compared to all other cell types. E. UMAP of prostate epithelial cells. Epithelial cells were subsetted and grouped into prostatic epithelial subtypes–luminal, basal, intermediate, and neuroendocrine cells. F. UMAP of epithelial cell marker gene expression showing cluster-specific enrichment. G. Dot plot of the top 10 upregulated genes of each epithelial cell cluster based on log2FC.

Uniform manifold approximation and projection (UMAP) based dimensionality reduction and unsupervised clustering revealed distinct cell populations that grouped primarily by cell type in the UMAP projection (Fig. 1C). Based on canonical markers, we annotated epithelial (*KRT8*, *KRT18*), endothelial (*KDR*, *FLT1*), pericyte (*RGS5*, *NOTCH3*), T cell (*CD3D*, *CD3E*), myeloid (*CD68*, *C1QB*), fibroblast (*DCN*, *LUM*), smooth muscle (*ACTG2*, *DES*), B cell (*CD79A*, *MS4A1*), and mast cell (*TPSAB1*, *TPSB2*) populations (Fig. S1). Differential gene expression confirmed cell-type specificity and reinforced these annotations (Fig. 1D).

To further explore the epithelial cells, we subclustered epithelial cell clusters and performed dimensionality reduction with PCA followed by UMAP. This revealed canonical luminal (*KLK3*, *ACPP*), basal (*KRT14*, *DST*), and rare neuroendocrine (*CHGA*, *CHGB*) subtypes (Fig. 1E, 1F). Strikingly, we identified a population of cells situated between basal and luminal clusters in the UMAP coordinate system. These cells often expressed *LTF* or *KRT13* (Fig. 1F) among a host of genes previously linked to prostate inflammatory atrophy (PIA) [13–17] and club/hillock cells [2], suggesting a potential intermediate epithelial state. Differential expression analysis highlighted unique gene signatures for these intermediate cells, including *LTF*, *LCN2*, *OLFM4*, *MMP7*, and *KRT13* (Fig. 1G).

### *WIF1*+ Basal Cells are Enriched in Human Prostate Transition Zone

Although prostate basal cells have traditionally been considered homogeneous across zones and patients, our analysis revealed substantial heterogeneity within this compartment. In initial UMAP projections, basal cell populations appeared separated by intermediate epithelial clusters (Fig. 1E), suggesting underlying transcriptional diversity and prompting further analysis.

To dissect this heterogeneity, we performed subclustering and identified four distinct basal subpopulations (Fig. 2A). The first, characterized by robust expression of canonical basal markers including *KRT5*, *KRT15*, and *KRT23*, was designated *KRT23*+ basal cells and exhibited a classical basal gene expression program (Fig. 2B, Fig. S2). The second, marked by *DDIT3*, *TXNIP*, *HES1*, and *MAFB* (Fig. 2B, Fig. S2), displayed a transcriptional program consistent with cellular stress responses and paligenosis [18,19], likely corresponding to the previously reported *PLCG2*+/c3 basal population [7]. The third, a minor population defined by *FOXI1* and vacuolar-type H+-ATPase subunits, resembled previously reported ionocyte-like basal cells (Fig. 2B, Fig. S2) [7].

**Figure 2.**
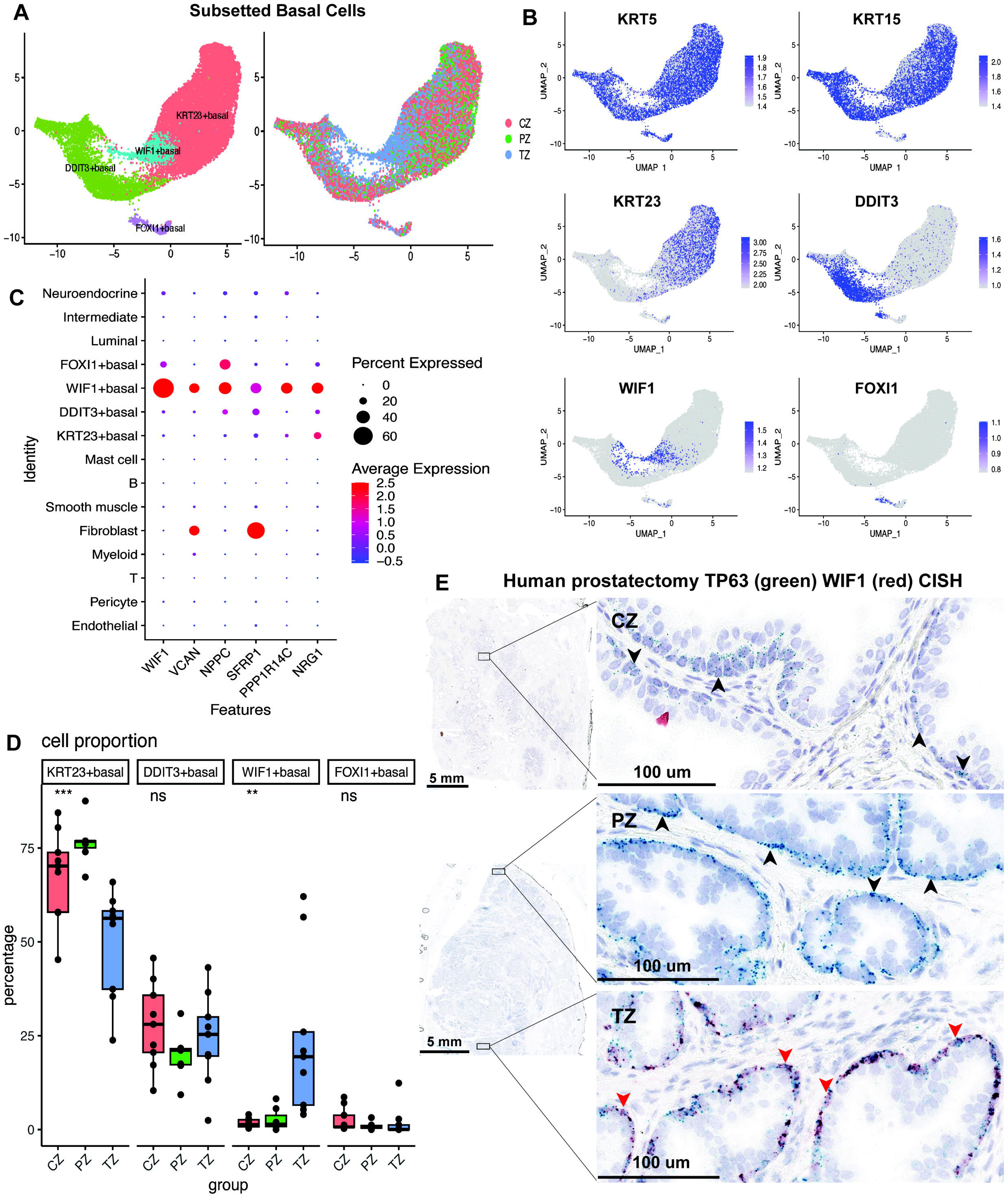
Zone-specific enrichment of prostatic basal subtypes. A. UMAPs of basal cell subset colored by cluster subtype (left) and zone (right). B. UMAPs of basal cell subsets with expression of marker genes indicated. *KRT5* and *KRT15*, canonical prostate basal cell markers were expressed in all basal cell subtypes, while *KRT23*, *DDIT3*, *WIF1*, and *FOXI1* showed subtype-specific expression. C. Dot plot showing expression of selected DEGs of *WIF1*+ basal cells across all prostate cell types. D. Box plots showing the proportion of basal cell subtype for each sample by prostate zones. Each dot represents a sample, and the box-whisker is colored by zone (CZ, PZ, TZ). Horizontal bars indicate the mean cell proportion of samples for each zone. P values were calculated using one-way ANOVA. ns: P > 0.05; *: P < 0.05; **: P < 0.01; ***: P < 0.001; ****: P < 0.0001. E. Representative examples of CISH staining of *TP63* (green), a known basal cell marker, and *WIF1* (red), a *WIF1*+ basal cell subtype marker in CZ, PZ, and TZ tissue samples from human prostatectomies. For 0.3x magnification, the scale bar indicates 5 mm. For 40x magnification, the scale bar indicates 100 μm. The red arrow indicates dual expression of *WIF1* and *TP63* in the TZ zone, while black arrows indicate the *TP63*+*/WIF1^-^* cells in CZ and PZ.

Notably, a fourth basal subpopulation exhibited high expression of *WIF1*, *VCAN*, *PPP1R14C*, *NPPC*, and *SFRP1* (Fig. 2B, Fig. S2), and closely resembled the previously reported Basal-5-VCAN subtype [6]. To further assess the specificity of these genes, we examined their expression across all prostate cell types. While *VCAN* and *SFRP1* were also expressed in fibroblasts, *WIF1* expression was exclusive and significantly enriched in this basal subtype (Fig. 2C). Based on the specificity, we designated this novel population as the *WIF1*+ basal cell subtype.

To determine whether basal subtypes displayed zonal preferences, we quantified their relative abundance across all samples. Strikingly, the *WIF1*+ basal subtype showed strong enrichment in the transition zone (TZ) (Fig. 2A, 2D).

To validate this striking zonal selectivity of the *WIF1*+ basal cell subtype, we performed chromogenic in situ hybridization (CISH) for *TP63*, a canonical basal marker, and *WIF1* on prostatectomy specimens from multiple zones. Co-localization of *TP63* and *WIF1* confirmed that *WIF1* expression is restricted to *TP63*+ basal epithelial cells, supporting the classification of *WIF1*+ cells as a distinct basal subtype. Notably, *TP63*+*WIF1*+ basal cells were detected exclusively in TZ, with nearly absent *WIF1* expression observed in basal cells of the PZ or CZ (Fig. 2E, Fig. S3).

We next examined whether this basal subtype exists in the mouse prostate. Intriguingly, *Wif1* was not expressed in any mouse basal epithelial cells. Instead, high expression was detected in a subset of fibroblasts (Fig. S4), corresponding to the ductal fibroblast population previously described by Joseph et al. [4] and subglandular fibroblast by our previous study, characterized by expression of *Rorb* [20]. These findings indicate that the *WIF1*+ basal cell subtype is specific to the human prostate.

### EMT, Stromal Remodeling, and Immune Modulation Pathways are Upregulated in *WIF1+* Basal Cells

Differential gene expression among the four basal subtypes revealed 104 genes uniquely upregulated in *WIF1*+ basal cells, compared to all other basal cell subtypes (Fig. 3A, Table S1). To explore the biological functions of this basal subtype, we performed pathway analysis using Hallmark and Gene Ontology Biological Process (GOBP) terms from MSigDB. *WIF1*+ basal cells were most significantly enriched for the Hallmark epithelial-mesenchymal transition (EMT) pathway, along with angiogenesis, androgen response, coagulation, and apical junction gene sets, suggesting a specialized role in tissue remodeling and zonal dynamics (Fig. 3B). GO analysis further implicated this subtype in prostate gland development and morphogenesis (Fig. 3C).

**Figure 3.**
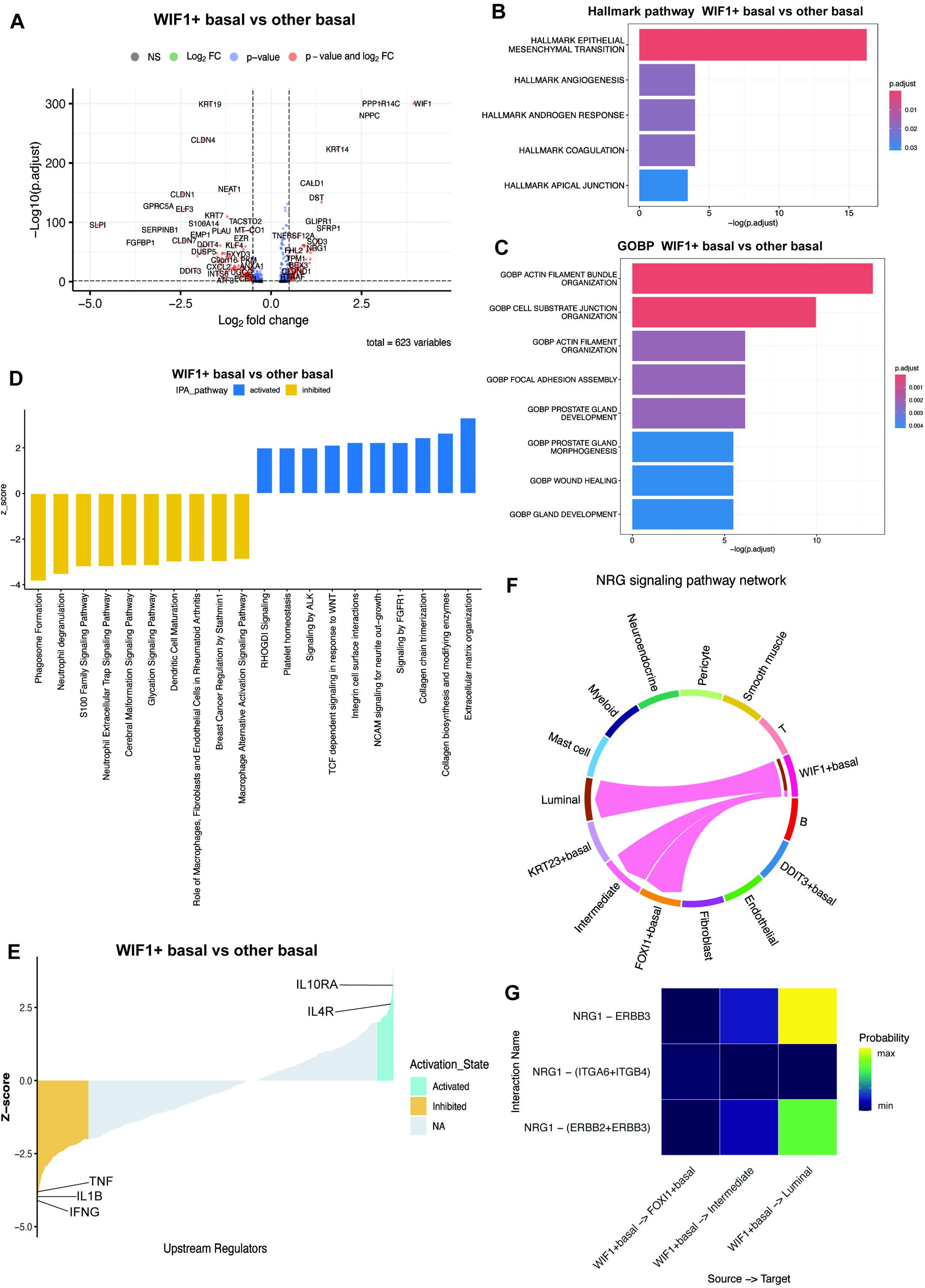
Gene expression pathway analysis of *WIF1*+ basal cell. A. Volcano plot showing DEGs identified between the *WIF1*+ basal cell and other basal cell subtypes. The x-axis indicates log2FC of gene expression. Log2FC > 0 denotes up-regulated genes in the *WIF1*+ basal cell compared to other basal cells. Log2FC < 0 denotes down-regulated genes in the *WIF1*+ basal cell compared to other basal cells. Red dots represent the significant DEGs with adjusted P < 0.05 and log2FC > 0.5. B. Barplot illustrating Hallmark pathway enrichment analysis of the genes up-regulated in the *WIF1*+ basal cell subtype relative to other basal cell subtypes. The DEGs were determined by adjusted P < 0.05 and log2FC > 0.5 The source of gene sets were derived from MSigDB (Molecular Signatures Database). Enriched pathways were performed by clusterProfiler and selected by adjusted p value < 0.05. C. Barplot illustrating GO-term enrichment analysis of the genes up-regulated in the *WIF1*+ basal cell subtype relative to other basal cell subtypes. The DEGs were determined by adjusted P < 0.05 and log2FC > 0.5. The source of gene sets were derived from MSigDB. Enriched pathways were performed by clusterProfiler and selected by adjusted p value < 0.05. D. IPA pathway analysis comparing WIF1+ basal cells to all the other basal cell subtypes. Pathways with z-scores > 0 are considered activated while z-scores < 0 are considered inhibited in WIF1+ basal cell subtype. The top 10 activated and inhibited Ingenuity Canonical Pathways are shown. E. IPA upstream regulator analysis comparing *WIF1*+ basal cells to all the other basal cell subtypes. Upstream regulators with z-scores > 2 are considered activated while z-scores < 2 are considered inhibited in *WIF1*+ basal cell subtype. Highlighted are *IL10RA* and *IL4R*, which are activated in the *WIF1*+ basal cell subtype compared to all the other basal cell subtypes, while *TNF*, *IL1B*, and *IFNG* are inhibited in the *WIF1*+ basal cell subtype compared to all the other basal cell subtypes. F. Chord diagram of the inferred *NRG* signaling pathway interactions between cell clusters. Edge width represents the interaction strength. A thicker edge line indicates a stronger signal. Edge direction was from the source cell to the target cell. G. Comparison of the key ligand-receptor pairs in *NRG* signaling pathway between different cell-cell communication groups. The color gradient represents the communication probability.

To complement these findings, we computationally inferred the activation and inhibition states of pathways and upstream regulators in *WIF1*+ basal cells compared to all other basal subtypes using Ingenuity Pathway Analysis (IPA). The top activated pathways included extracellular matrix organization, collagen biosynthesis and modifying enzymes, collagen chain trimerization (Fig. 3D, Table S2), suggesting a potential role for *WIF1*+ basal cells in extracellular matrix and basement membrane remodeling and glandular tissue architecture support. Based on these observations, we speculated that these *WIF1*+ basal cells may have activated tissue-structural programs.

In contrast, immune-associated pathways such as phagosome formation, neutrophil degranulation, and S100 family signaling were predicted to be inhibited in *WIF1*+ basal cells. Likewise, The top inhibited upstream regulators of *WIF1*+ basal cells included *IFNG*, *IL1B*, *NFKB*, and *TNF*, while the top activated regulators included several anti-inflammatory cytokines, such as *IL10RA*, *IL4R* (Fig. 3E, Table S3). Collectively, these analyses suggested an immunomodulatory role for *WIF1*+ basal cells within the transition zone microenvironment. Based on these observations, we speculated that these *WIF1*+ basal cells may have inhibited immunogenic programs.

To investigate potential intercellular communication, we applied ligand-receptor interaction analysis across all cell types in the human prostate. Notably, *NRG* (neuregulin) signaling was inferred to occur exclusively from *WIF1*+ basal cells to other epithelial populations (Fig. 3F). A chord diagram visualized the directionality and strength of the interaction, revealing that *WIF1*+ basal cells act as a unique ligand source, while luminal, intermediate, and *FOXI1*+ basal cells serve as target populations. Among these, luminal cells exhibited the strongest inferred interaction. Looking into the ligand-receptor pair in *NRG* signaling, *NRG1* produced by *WIF1*+ basal cells was predicted to interact with *ERBB2*/*ERBB3* receptors on luminal cells (Fig. 3G). Given that *NRG*-*ERBB* signaling plays a key role in epithelial morphogenesis and homeostasis [21], these findings suggest that *WIF1*+ basal cells may serve as the regulator of epithelial maintenance and differentiation within the transition zone niche.

### *WIF1*+ Basal Cells Are Enriched in BPH

Given the enrichment of *WIF1*+ basal cells in the transition zone and involvement in pathways such as EMT [12], androgen response [8], and stromal remodeling [10], we hypothesized that this basal subtype may be involved in the pathogenesis of benign prostatic hyperplasia (BPH).

To test this, we analyzed a previously published single-cell RNA-seq dataset [4], which included both BPH tissues and normal adult prostates. Dimensionality reduction and clustering of this dataset revealed clear segregation by canonical cell types (Fig. 4A). Upon subsetting basal epithelial cells, we identified the same four transcriptionally distinct basal subtypes described in our dataset, marked by *KRT23*, *DDIT3*, *WIF1*, and *FOXI1*, respectively (Fig. 4B-C). The *WIF1*+ basal cells from this dataset exhibited a highly similar gene signature to that in our prostatectomy samples, including expression of *WIF1*, *VCAN*, *NPPC*, and *PPP1R14C* (Fig. S5), strongly supporting the identity of this population as the same cell type.

**Figure 4.**
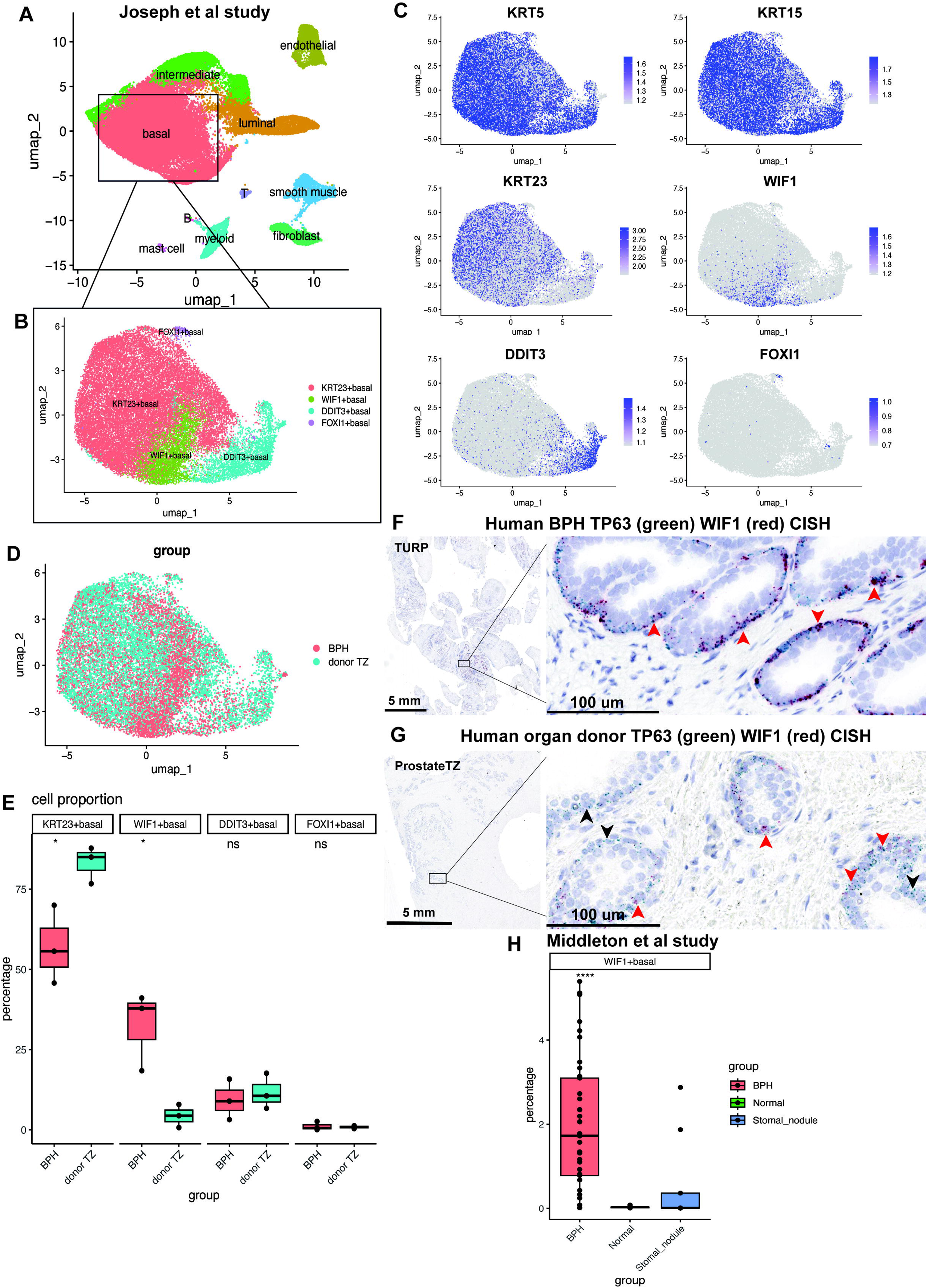
*WIF1*+ basal cells are enriched in BPH. A. UMAP and clustering analysis of previously published scRNA-seq dataset of BPH and normal prostate from organ donor by Joseph et al. [4], showed cell clustering by known cell types in the prostate. B. UMAP of four basal cell clusters, labeled with dominantly expressed marker genes for each subtype. C. UMAP of each marker gene showed basal cell subtype-specific expression. *KRT5* and *KRT15*, known prostatic basal cell markers, were positive for all basal cell subtypes, while subtype-specific marker genes (*KRT23, DDIT3, WIF1,* and *FOXI1*) were expressed in corresponding basal cell subclusters. D. UMAP of basal cell subset labeled by groups (BPH, donor TZ). E. Box plots showing the proportion of basal cell subtypes for each sample. Each dot represents a sample and the box-whiskers are colored by groups (BPH, donor TZ). Horizontal bars indicate the mean cell proportion of samples for each group. P values were calculated using a two-sided *t*-test. ns: P > 0.05; *: P < 0.05; **: P < 0.01; ***: P < 0.001; ****: P < 0.0001. F. Representative example of *TP63* (green) and *WIF1* (red) CISH staining in BPH tissue of human TURP (Transurethral Resection of the Prostate) samples. For 0.3x magnification, the scale bar indicates 5 mm. For 40x magnification, the scale bar indicates 100 μm. The red arrow indicates dual expression of *WIF1* and *TP63* in the BPH patients. G. Representative example of *TP63* (green) and *WIF1* (red) CISH staining in the TZ gland of human organ donor samples. For 0.3x magnification, the scale bar indicates 5 mm. For 40x magnification, the scale bar indicates 100 μm. The red arrow indicates dual expression of *WIF1* and *TP63* in the TZ zone of prostate donor. The black arrow indicates a *TP63*-expressing cell lacking *WIF1* in the TZ of the prostate donor. H. Cell type deconvolution analysis by RCTD (Robust Cell Type Decomposition) of bulk RNA-seq dataset from BPH, normal prostate, and BPH stromal nodules by Middleton et al. [22]. Proportion of *WIF1*+ basal cell type was estimated for each sample. Each dot represents one bulk RNA-seq sample. Box-whiskers are colored by group (BPH, Normal, BPH stromal nodule). Horizontal bars indicate mean cell proportions per group. P values were calculated using one-way ANOVA. ns: P > 0.05; *: P < 0.05; **: P < 0.01; ***: P < 0.001; ****: P < 0.0001.

We next quantified the relative abundance of basal subtypes across BPH and normal TZ samples from organ donors. The proportion of *WIF1*+ basal cells was significantly elevated in BPH tissues compared to donor TZ (Fig. 4D-E).

Next, we analyzed the BPH stromal nodule dataset from the same study [4]. Using canonical marker genes, we identified major prostate cell types within the stromal nodules. In this dataset, epithelial cells represented only a small fraction compared to stromal populations such as fibroblasts and smooth muscle cells. Notably, within the epithelial compartment, a significant fraction of basal cells expressed *WIF1* (Fig. S6).

To confirm this observation at the tissue level, we performed CISH on BPH samples obtained via transurethral resection of the prostate (TURP), alongside TZ tissues from young organ donors from JHU. Co-staining for *TP63* and *WIF1* confirmed the identity of *WIF1*+ basal cells, and these cells were significantly abundant in BPH specimens (Fig. 4F-G, Fig. S7).

To further validate this enrichment, we analyzed previously published bulk RNA-seq data from BPH and normal prostate tissues [22]. Notably, several *WIF1*+ basal cell specific genes, including *WIF1*, *NPPC*, and *VCAN*, were among the most significantly upregulated DEGs in BPH compared with normal prostate (Fig. S8). Using our scRNA-seq dataset as a reference, cell type deconvolution analysis further confirmed a significant enrichment of *WIF1*+ basal cells in BPH samples relative to both normal prostate and BPH stromal nodule samples (Fig. 4H, Fig. S8).

### *WIF1*+ Basal-Specific Genes are Upregulated in BPH

To determine whether *WIF1*+ basal cells undergo transcriptional changes in BPH, we compared their transcriptomes in BPH and normal TZ from organ donors using the previously published single cell sequencing dataset [4]. A large number of ribosomal genes, including *RPS10*, *RPS29*, *RPL39*, *RPL12*, were significantly upregulated in *WIF1*+ basal cells from BPH compared to those from normal TZ (Fig. 5A, Table S4). Gene ontology (GO) analysis of the differentially expressed genes (DEGs) revealed strong enrichment of pathways involved in ribosomal biogenesis (Fig. 5B), suggesting elevated biosynthetic activity in these cells during BPH development. Oxidative stress response pathways and cell adhesion pathways were also upregulated.

**Figure 5.**
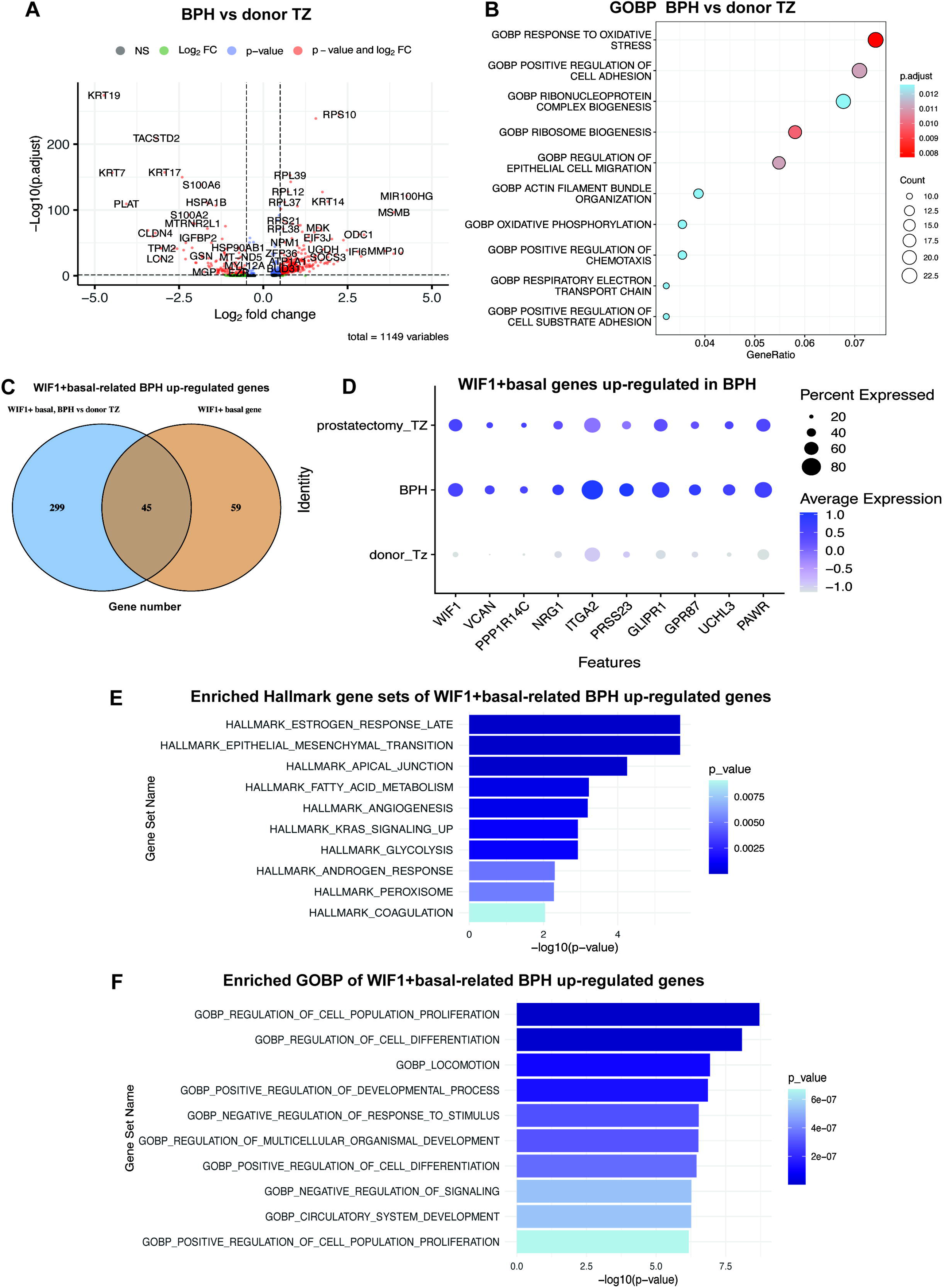
*WIF1*+ basal-specific genes are upregulated in BPH. A. Further analysis of previously published scRNA-seq dataset of BPH and normal prostate from organ donors by Joseph et al. [4]. Volcano plot showing DEGs identified between the *WIF1*+ basal cell subtype in BPH group and *WIF1*+ basal cell subtype in TZ of the organ donor group. The x-axis indicates log2FC of gene expression. Log2FC > 0 denotes up-regulated genes in BPH compared to donor TZ. Log2FC < 0 denotes down-regulated genes in BPH compared to donor TZ. Red dots represent the significant DEGs with adjusted P < 0.05 and log2FC > 0.5. B. Dotplot illustrating GO-term enrichment analysis of the genes up-regulated in BPH compared to donor TZ within the *WIF1*+ basal cell subtype. The top 10 pathways with adjusted P value < 0.05 were selected for visualization. The count indicates the number of DEGs associated with each gene set. The GeneRatio indicates the number of DEGs associated with each gene set divided by the total number of DEGs. C. Venn diagram depicting the number of *WIF1*+ basal-related BPH up-regulated genes, overlapping from two comparisons-BPH versus donor TZ within the *WIF1*+ basal cell subtype, and *WIF1*+ basal versus other basal subtypes. D. Dot plot showing the expression of representative *WIF1*+ basal related BPH up-regulated genes in *WIF1*+ basal cell subtype of prostatectomy TZ, BPH, and donor TZ. E. Barplot illustrating Hallmark pathway enrichment analysis of *WIF1*+ basal-related BPH up-regulated genes. The MSigDB online platform (https://www.gsea-msigdb.org/gsea/msigdb/index.jsp) was used to identify Hallmark gene sets enriched for the *WIF1*+ basal genes upregulated in BPH. The top 10 pathways with a false discovery rate (FDR) q-value < 0.05 were selected for visualization. F. Barplot illustrating GO-term enrichment analysis of *WIF1*+ basal-related BPH up-regulated genes. The MSigDB online platform (https://www.gsea-msigdb.org/gsea/msigdb/index.jsp) was used to identify GOBP enriched for the *WIF1*+ basal genes upregulated in BPH. The top 10 pathways with a false discovery rate (FDR) q-value < 0.05 were selected for visualization.

To pinpoint genes associated with both cell identity and disease, we intersected DEGs from two comparisons: 1) *WIF1*+ basal versus other basal subtypes (current study), and 2) BPH versus donor TZ from *WIF1*+ basal cell (derived from Joseph et al. [4]). This yielded 45 overlapping upregulated genes, that we refer to as “*WIF1*+ basal-related BPH up-regulated genes” (Fig. 5C, Table S5). Among these, the most significantly elevated genes included *WIF1*, *VCAN*, *PPP1R14C*, *NRG1*, *ITGA2*, *PRSS23*, *GLIPR1*, *GPR87*, *UCHL3*, and *PAWR* (Fig. 5D). Several of these genes are key components of the *WIF1*+ basal cell signature and were also upregulated in BPH, supporting their potential relevance in disease pathology. For example, upregulation of *WIF1*, a known WNT signaling inhibitor [23], and *VCAN*, a major extracellular matrix (ECM) proteoglycan, suggested a role in modulating epithelial-stromal signaling and extracellular matrix remodeling [24]. *PPP1R14C*, a regulatory inhibitor of protein phosphatase 1, may contribute to cytoskeletal or contractile regulation [25]. Notably, *NRG1*, which was predicted to signal from *WIF1*+ basal cells to *ERBB2*/*ERBB3* receptors on neighboring epithelial cells, was also elevated in BPH, suggesting a potential paracrine signaling role. Other genes, such as *PRSS23* (a serine protease that cleaves peptide bonds in proteins) [26], *GPR87* (encodes a G protein-coupled receptor), found to contribute to fibrosis [27], and *UCHL3* (ubiquitin C-terminal hydrolase L3), a protein that plays a crucial role in cellular processes by removing ubiquitin from other proteins) [28], further suggested involvement in cell proliferation and epithelial-stromal remodeling. Notably, these genes displayed a graded expression pattern across conditions, with the highest levels observed in BPH, the lowest in the donor TZ, and intermediate levels in TZ samples from prostatectomy (Fig. 5D), many of which showed evidence of nodular hyperplasia consistent with BPH.

Pathway enrichment analysis on *WIF1*+ basal-related BPH up-regulated genes revealed activation of Epithelial-mesenchymal transition (EMT), along with Estrogen response, Apical junction, Angiogenesis pathways (Fig. 5E), indicating potential Epithelial-mesenchymal transition, hormonal sensitivity, and stromal activity. GOBP terms such as cell proliferation, cell differentiation, and cell locomotion were also significantly enriched (Fig. 5F). Taken together, we speculated that TZ enriched *WIF1*+ basal cells may play an active role in BPH pathogenesis by participating in remodeling of the prostate epithelium and stroma.

## DISCUSSION

Human prostate epithelium comprises four principal cell types: luminal, basal, intermediate, and a small population of neuroendocrine epithelial cells. Among these, subtypes within the basal cell compartment have been rarely characterized in the literature. In this study, we identified *WIF1*+ basal cells as a transcriptionally and anatomically distinct subpopulation compared to other basal epithelial cells. These cells display a unique gene expression profile enriched for *WIF1*, *VCAN*, *PPP1R14C*, and *NRG1*, consistent with the Basal-VCAN cell type described by Hu et al [6]. Gene Ontology analysis revealed that *WIF1*+ basal cells are enriched in pathways related to gland development and morphogenesis, further supporting their correspondence to the Basal-VCAN population. While Hu et al. reported that this cell type is predominantly located in both the peripheral and transition zones [6], our scRNA-seq data and CISH staining demonstrated that *WIF1*+ basal cells are markedly enriched in the transition zone (TZ) of the human prostate.

Among the upregulated genes, *WIF1* functions as a Wnt signaling antagonist. The Wnt signaling pathway is a critical regulatory network involved in embryonic development, cell proliferation, and carcinogenesis [29–35]. Aberrant Wnt activation has been implicated in prostate cancer progression [36,37]. Studies have shown that WIF1 is down-regulated in prostate cancer [38]. Restoration of WIF1 expression in prostate cancer cells suppresses tumor growth, reduces migration and invasion, and reverses epithelial-to-mesenchymal transition [39].

In addition to *WIF1*, two other classes of secreted Wnt antagonists have been described: the secreted Frizzled-related protein (sFRP) family [40] and the Dickkopf (Dkk) family [41]. Notably, in our dataset, both *SFRP1* and *DKK3* were also highly expressed in *WIF1*+ basal cells (Table S1). The combined high expression of *WIF1*, *SFRP1*, and *DKK3* in the basal layer of the TZ may establish a localized Wnt signaling gradient, limiting the ability of stromal Wnt ligands to activate epithelial receptors [42].

Furthermore, IPA analyses revealed that inflammation-related pathways were suppressed in *WIF1*+ basal cells relative to other basal populations. Together, these findings lead us to speculate that *WIF1*+ basal cells in the TZ help create an epithelial microenvironment characterized by attenuated WNT activation and reduced inflammation, potentially preserving epithelial homeostasis and contributing to the relatively low incidence of prostate cancer initiation in the TZ. This hypothesis warrants further mechanistic investigation.

Benign prostatic hyperplasia (BPH) is a highly prevalent age-associated condition characterized by epithelial and stromal hyperplasia, predominantly in the transition zone (TZ) of the prostate [43]. Analysis of publicly available datasets, combined with CISH staining of BPH tissues from TURP patients, revealed an enrichment of *WIF1*+ basal cells in BPH. Furthermore, the gene signature of *WIF1*+ basal cells was distinctly altered in BPH compared to donor TZ, suggesting that this TZ-enriched population may contribute to BPH pathogenesis.

Among the top differentially expressed genes in *WIF1*+ basal cells, *VCAN*, encoding the extracellular matrix (ECM) component versican, points to active involvement in ECM remodeling. Versican has been shown to promote fibroblast activation and myofibroblast-like phenotypes [44,45], potentially driving the fibromuscular expansion characteristic of BPH. Pathway enrichment analysis identified epithelial mesenchymal transition (EMT) as the most significant hallmark pathway in *WIF1*+ basal cells. EMT, a process by which epithelial cells acquire mesenchymal features, underlies tissue plasticity, regeneration, and fibrosis [46]. Its activation in *WIF1*+ basal cells aligns with previous evidence implicating EMT in BPH etiology [12].

*NRG1*, another gene differentially expressed in *WIF1*+ basal cells and enriched in BPH, is known to activate *ERBB2/3* receptors on neighboring epithelial cells, thereby stimulating proliferation and survival [47]. Its role in promoting epithelial regeneration has been demonstrated in the intestinal stem cell niche [48]. Our cell-cell communication analysis inferred that *NRG* signaling occurs exclusively between *WIF1*+ basal cells and other epithelial cells, particularly luminal cells. We propose that *WIF1*+ basal cells may secrete NRG1 to regulate adjacent luminal epithelium, establishing a paracrine loop that promotes epithelial development in BPH. Interestingly, previous studies have reported NRG1 secretion by fibroblasts, which then interact with EGFR and ERBB3 receptors on epithelial cells [4,6].

Our scRNA-seq data indicated that *Wif1* is not expressed by basal cells in the mouse prostate but is highly expressed in a subset of fibroblasts, termed *Rorb*+ subglandular fibroblasts [20], suggesting that *WIF1*+ basal cells represent a human-specific cell population. Notably, *WIF1*, *VCAN*, and *NRG1* genes reported to be associated with fibroblasts, are highly expressed in *WIF1*+ basal cells, raising the possibility that these cells exhibit mesenchymal-like characteristics. In a previous study, Alonso-Magdalena et al. concluded that BPH stroma can arise from epithelium through epithelial-mesenchymal transition (EMT), a process in which epithelial cells lose their polarity and basement membrane attachment while acquiring mesenchymal features [12]. Whether *WIF1*+ basal cells enriched in TZ and BPH represent an intermediate state between basal epithelial and mesenchymal cells warrants further investigation.

The identification of *WIF1*+ basal cells may also have implications for zone-specific susceptibility. *WIF1*+ basal cells in TZ may provide a microenvironment that supports epithelial-mesenchymal transition, proliferation, extracellular matrix organization, and angiogenesis, whereas the PZ and CZ may lack this *WIF1*+ basal cell-mediated function. This could partly explain why BPH preferentially develops in the TZ.

A key limitation of our study is the absence of direct functional experiments to validate the inferred roles of *WIF1*+ basal cells in BPH pathogenesis. Although our single-cell transcriptomic analysis, pathway enrichment results, and histological validation collectively support their involvement in EMT activation, extracellular matrix remodeling, cell proliferation, and modulation of WNT signaling, these conclusions remain correlative. Without functional assays, such as lineage tracing, in vitro co-culture systems, or in vivo animal models, it remains uncertain whether *WIF1*+ basal cells are causal drivers or secondary responders in the observed tissue remodeling. Future studies integrating targeted manipulation of *WIF1* expression or depletion of this basal cell population will be essential to establish causality and delineate their precise mechanistic contributions to BPH development. Spatial transcriptomic approaches will also be crucial for mapping their interactions with stromal, immune, and vascular components in situ.

## MATERIALS AND METHODS

### Human prostatectomy tissue punches

Prostate tissue specimens were collected from men diagnosed with primary prostate cancer undergoing radical prostatectomy at Johns Hopkins University and consented under IRB approved protocols. Prostatectomies were sectioned fresh from apex to base, and fresh tissue samples were collected using an 8 mm punch biopsy tool from each zone of the prostate. A quick HE staining was performed to confirm no cancer lesion included in these punch areas. Freshly collected tissue punches were processed for scRNA-seq [13].

### Dissociation of human prostate tissues

Dissected tissues were minced with razor blades and digested in 0.25% Trypsin-EDTA (Gibco 25200-072) for 10 min at 37 °C, followed by incubation for 2.5 hours at 37 °C with gentle agitation in DMEM containing 10% FBS, 1 mg/mL Collagenase Type I (Gibco 17100-017), and 0.1 mg/mL of DNase I (Roche 10104159001). Digested tissues were centrifuged at 400 x g for 5 minutes, washed with HBSS, and further incubated in 0.25% Trypsin-EDTA for 10 min at 37 °C. Cells were suspended in DMEM containing 10% FBS and 0.4 mg/mL of DNase I and filtered through a 40 μm cell strainer.

### Single-cell RNA-sequencing and data pre-processing

Libraries for scRNA-seq were prepared using the 10x Genomics Chromium Single Cell 3’ Library and Gel bead Kit V2 (CG00052_RevF) according to the manufacturer’s protocol for each human prostatectomy tissue punch. The cDNA libraries were sequenced (150 bp paired-end) on the Illumina HiSeqX platform. Each sequenced library was demultiplexed to FASTQ files using Cell Ranger (10x Genomics). Cell Ranger (version 3.2.0) count pipeline was used to align reads to the GRCh38 transcriptome and create a gene-by-cell count matrix. The resulting gene expression matrix was loaded into Seurat (v4.0.4) in R. Low-quality cells were filtered out if genes were below 500, as well as if mitochondrial genes were >20%.

### Data Integration and cell type identification

We applied the “anchor-based” integration method to assemble multiple samples into an integrated dataset, following the Seurat integration workflow [49]. After running principal component analysis (PCA), FindNeighbors and FindClusters functions, we performed nonlinear dimensionality reduction with the RunUMAP function to obtain a two-dimensional representation. Differential gene expression analysis of previously characterized cell type-specific genes was used to identify the cell type for each cluster. A small cluster of cells expressing biomarkers from more than one cell type (epithelial, stromal, and immune) were considered as doublets and removed from downstream analysis.

### Differential gene expression analysis

The FindMarker function in the Seurat package was performed to identify differentially expressed genes (DEGs) among cell clusters or samples. Statistical significance was assessed using the Wilcoxon rank-sum test, and P-values were adjusted for multiple testing using the Benjamini-Hochberg method. Genes with log₂(fold change) > 0.5 and adjusted P < 0.05 were considered significant DEGs.

### Pathway enrichment analysis

Gene sets from the Molecular Signatures Database (MSigDB), including the Hallmark and Gene Ontology Biological Process (GOBP) collections, were imported using the R package msigdbr [50]. Pathway enrichment analysis of significantly differentially expressed genes (DEGs) was performed using clusterProfiler with the hypergeometric test [51]. Significantly enriched pathways were identified based on adjusted p-values. For analyses with a limited number of DEGs, specifically *WIF1*+ basal genes upregulated in BPH and identified by intersecting two comparisons, pathway enrichment was performed using Gene Set Enrichment Analysis (GSEA) via the MSigDB online platform (https://www.gsea-msigdb.org/gsea/msigdb/index.jsp). The top 10 pathways with a false discovery rate (FDR) q-value < 0.05 were selected for visualization.

### Ingenuity Pathway Analysis (IPA)

Upstream regulators that are likely to mediate the observed gene expression differences across basal cell clusters were analyzed using Ingenuity Pathway Analysis (IPA) [52]. Biological networks constructed from known interactions in the published literature were used to infer upstream molecular regulators and pathways. Using observed differential gene expression, a z-score was derived from predicted up or down-regulation of relevant genes in a pathway. Seurat was used to perform differential gene expression analyses comparing *WIF1*+ basal to other basal cell clusters. The subsequent IPA analysis was implemented with differentially expressed genes with at least abs(log2FC) > 1 and a false discovery rate adjusted-p-value < 0.01. The predicted pathways and upstream regulators were plotted with their associated z-scores.

### Chromogenic in situ hybridization (CISH)

To prepare FFPE tissue for CISH staining, slides were incubated for 30 minutes at 60 °C and deparaffinized by incubating slides at room temperature (RT) for 10 minutes in xylene twice, and then subsequently incubated in 100% ethanol twice and finally left to air dry. Hydrogen peroxide solution was added to the slides for 10 minutes at RT. Slides were steamed in 1 x RNAscope Target retrieval reagent at 100 °C for 18 minutes, followed by protease plus digestion for 30 minutes at 40 °C to allow target accessibility. The following probes were used for CISH staining, including Hs-*TP63* (RNAscope, Cat No. 601891) and Hs-*WIF1* (RNAscope, Cat No.429391-C2). Probes were added to slides and incubated in the HybEZ TM Oven for 2 hours at 40 °C, followed by signal amplification and detection assay according to the manufacturer’s protocol. The C1 probe signal was detected with Green color and C2 probe signal was detected with Red color. Slides were counter-stained with 50% Gill’s Hematoxylin for 30 seconds, baked for 15 min at 60 °C, coverslipped with VectaMount AQ Aqueous mounting medium, and then scanned with VENTANA DP200 (ROCHE).

### Cell communication network analysis

CellChat [53] was used to explore the cell-cell communication between cell types. First, the normalized gene expression data was converted into a CellChat object. A ligand-receptor interaction database was used to predict signaling networks based on gene expression patterns. Next, communication probabilities were computed at the signaling pathway level by aggregating the probabilities of all ligand-receptor interactions associated with each pathway. We filtered out communications if there were fewer than 10 cells in the cell group.

### Single-cell RNA-sequencing of mouse prostate

Mouse prostate tissues and scRNA-seq data were processed as previously described [20].

### Publicly available datasets

We additionally processed scRNA-seq data of human normal and BPH tissues by Joseph DB et al. [4]. Three datasets of BPH glandular samples (GSM5252126, GSM5252128, GSM5252130), BPH stromal samples (GSM5252127, GSM5252129, GSM5252131), and six datasets of young organ donor samples (GSM5252457, GSM5252459, GSM5252461, GSM5252458, GSM5252460, GSM5252462), were obtained from the GEO database under accession code GSE172357. We processed the data using a similar strategy described above to filter low-quality cells, integrate and cluster cells. Then the cell type was identified, and differential gene expression analysis was performed among cell types.

Bulk RNA-seq dataset of BPH, BPH stromal nodules, and normal prostate from Middleton et al. [22] were obtained from the dbGAP database, with accession number phs0016 98.v1.p1. RCTD [54] was applied to deconvolute cell types in the bulk RNA-seq data using our human prostatectomy single cell RNA-seq dataset as reference.

The mouse prostate data were from Graham MK et al. [20], and are deposited in the NCBI GEO database under accession code GSE228945.

## AUTHOR CONTRIBUTIONS

Conceptualization, R.W., A.M.D., and S.Y.; Methodology, R.W., Q.Z., A.M.D., and S.Y.; Validation, Q.Z., A.M.D., and S.Y.; Formal Analysis, R.W., S.Y.; Investigation, R.W., Q.Z., M.K.G., A.V., J.L., J.G., T.J., N.C., Y.Y., W.G.N., A.M.D., and S.Y.; Resources, Q.Z., T.J., W.G.N., A.M.D., and S.Y.; Data Curation, R.W., A.G., Y.Z., K.S., J.M., A.S., D.H., and S.Y.; Writing-Original Draft, R.W., and S.Y.; Supervision, W.G.N., A.M.D., and S.Y.; Funding Acquisition, W.G.N., A.M.D., and S.Y.; Manuscript editing and approval: all authors.

## ACKNOWLEDGEMENTS

We thank the members of the Sidney Kimmel Comprehensive Cancer Center’s Experimental and Computational Genomics Core, supported by Cancer Center Support Grant P30CA006973 (W.G.N., S.Y.), for support with the single-cell sequencing studies and data pre-processing and analysis. This work was supported in part by NIH/NCI grants P50CA058236 (W.G.N., S.Y., A.M.D.), U01CA196390 (A.M.D., S.Y.), P01CA247886 (S.Y.), U54CA274370 (A.M.D., S.Y.), P50CA180995 (C.E.P. awarded to M.K.G.), and by the Prostate Cancer Foundation (S.Y.), The Allegheny Health Network Johns Hopkins Pilot Project Grant (S.Y.), The Patrick C. Walsh Fund (S.Y.), The Irving Hansen Foundation (S.Y.), The Commonwealth Foundation (S.Y.), and the Maryland Cigarette Restitution Fund (S.Y.).

## SUPPLEMENTARY FIGURES

**Supplementary Figure 1.**
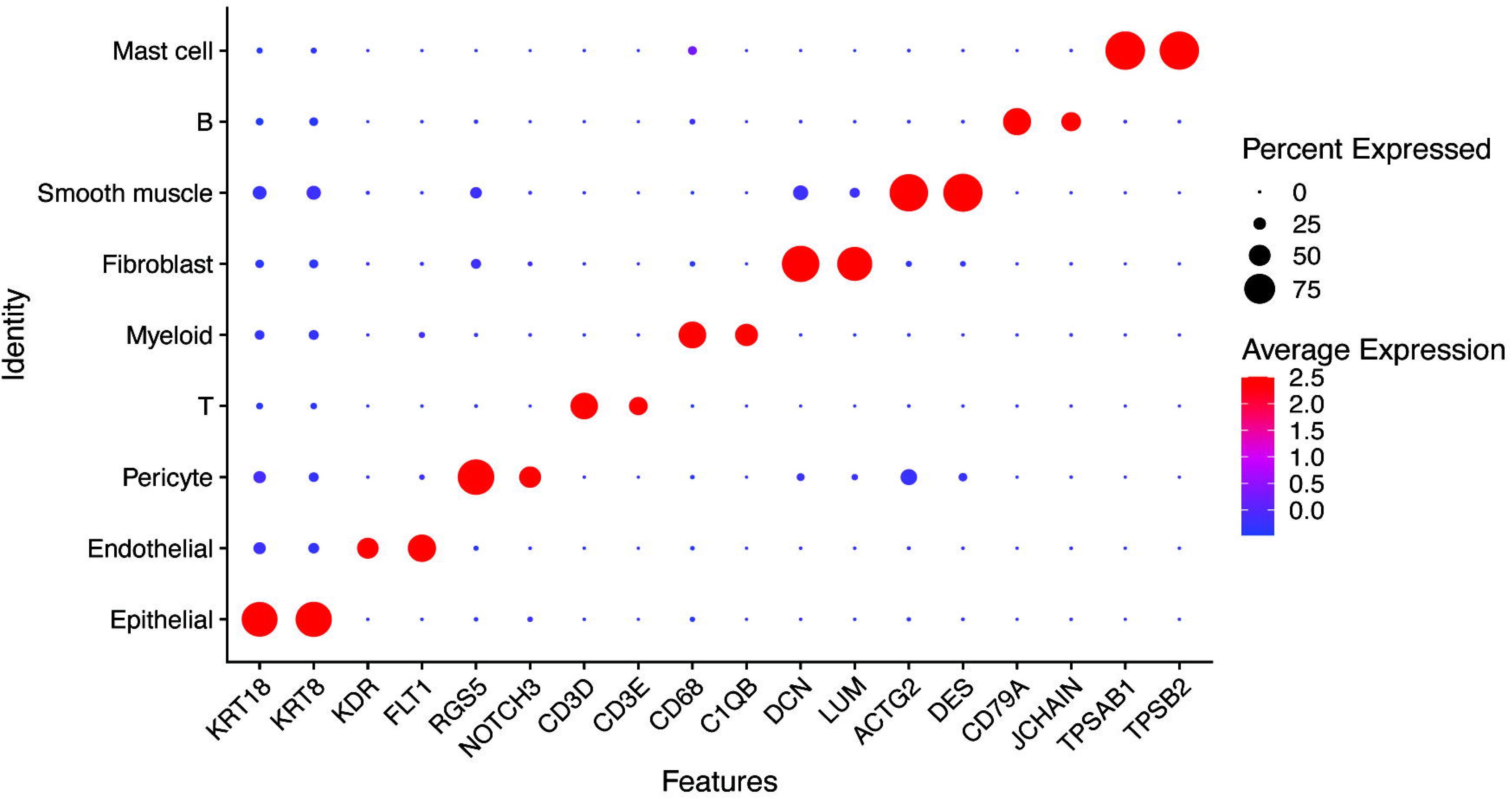
Dotplot of expression of prostate marker genes in human prostatectomy samples.

**Supplementary Figure 2.**
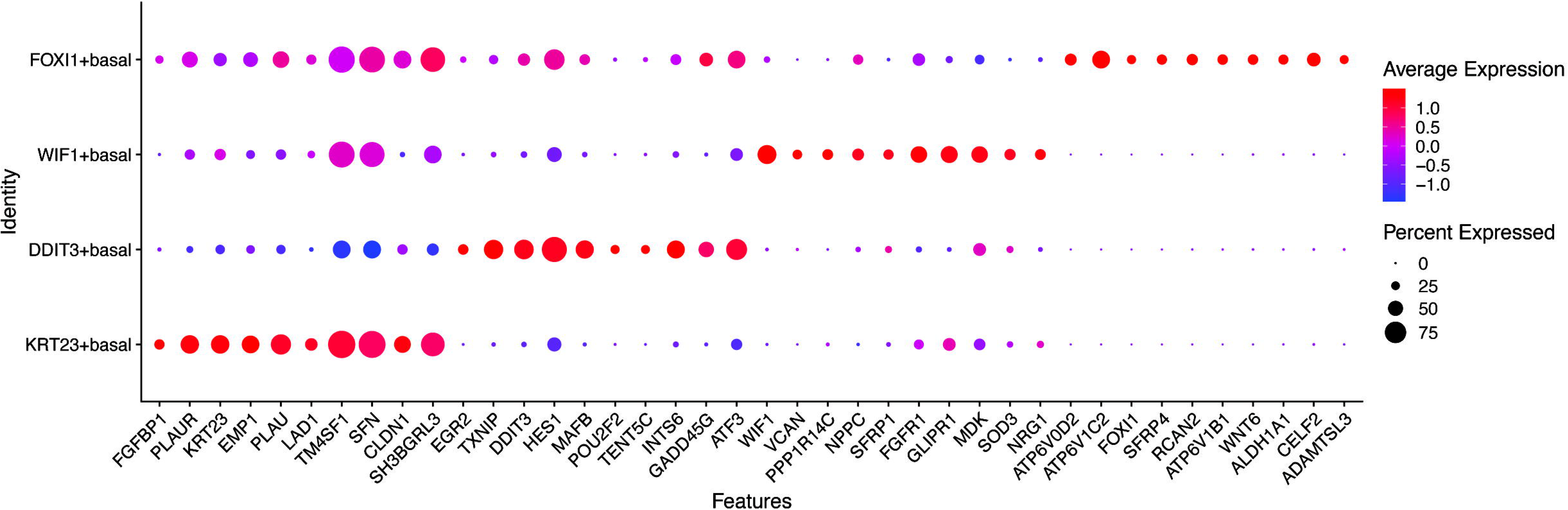
Dotplot of top 10 differentially expressed genes by adjusted p-value and log fold change for different basal cell subtypes of human prostatectomy samples.

**Supplementary Figure 3.**
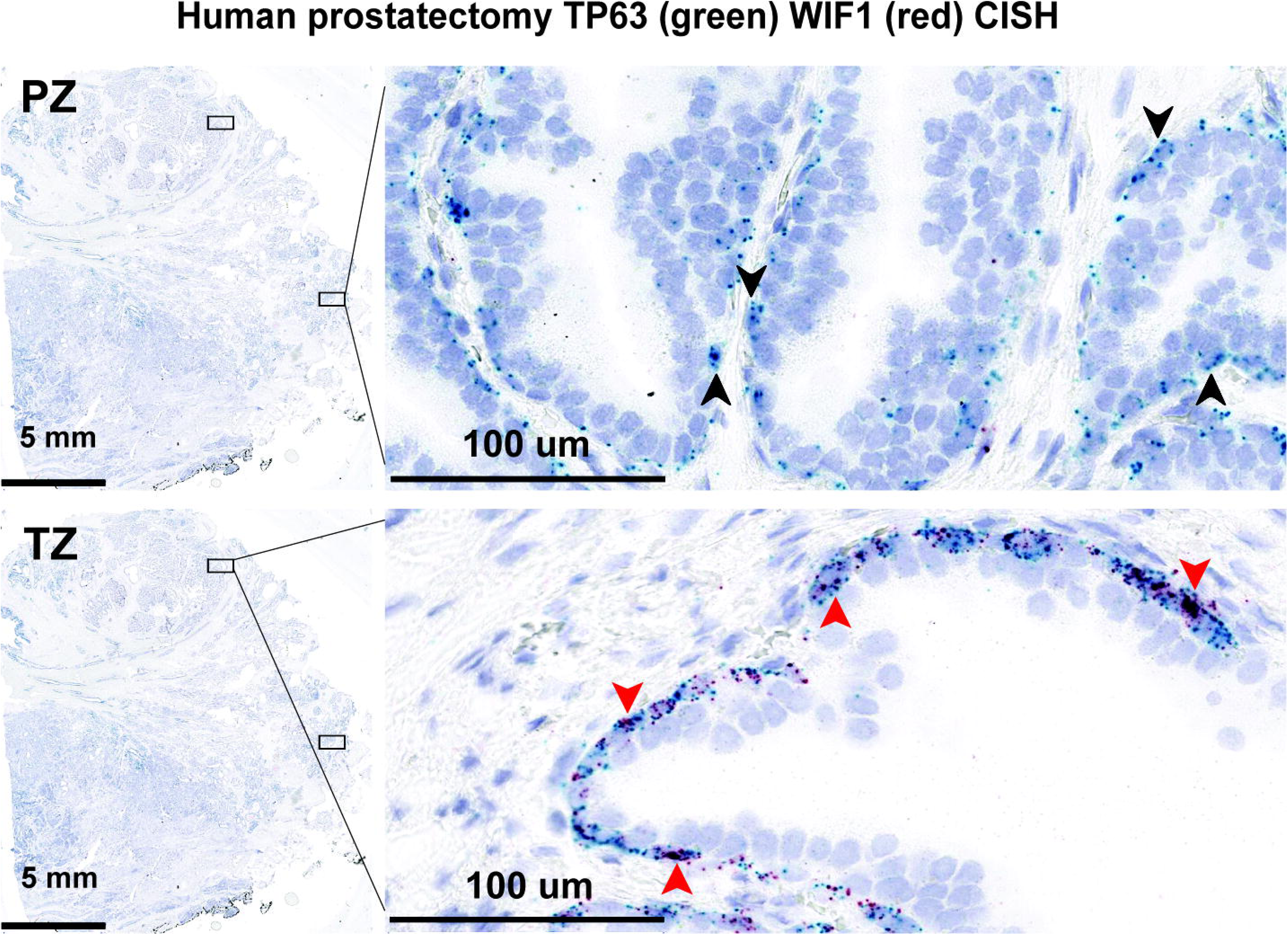
*WIF1*+ basal Cell is Enriched in Human Prostate Transition Zone. Representative examples of CISH staining of *TP63* (green), a known basal cell marker, and *WIF1* (red), a *WIF1*+ basal cell subtype marker, in PZ and TZ gland of human prostatectomy tissue samples. For 0.3x magnification, the scale bar indicates 5 mm. For 40x magnification, scale bar indicates 100 μm. The red arrow indicates dual expression of *WIF1* and *TP63* in the TZ zone, while lack of dual positive cells in PZ is indicated by black arrows.

**Supplementary Fig. 4.**
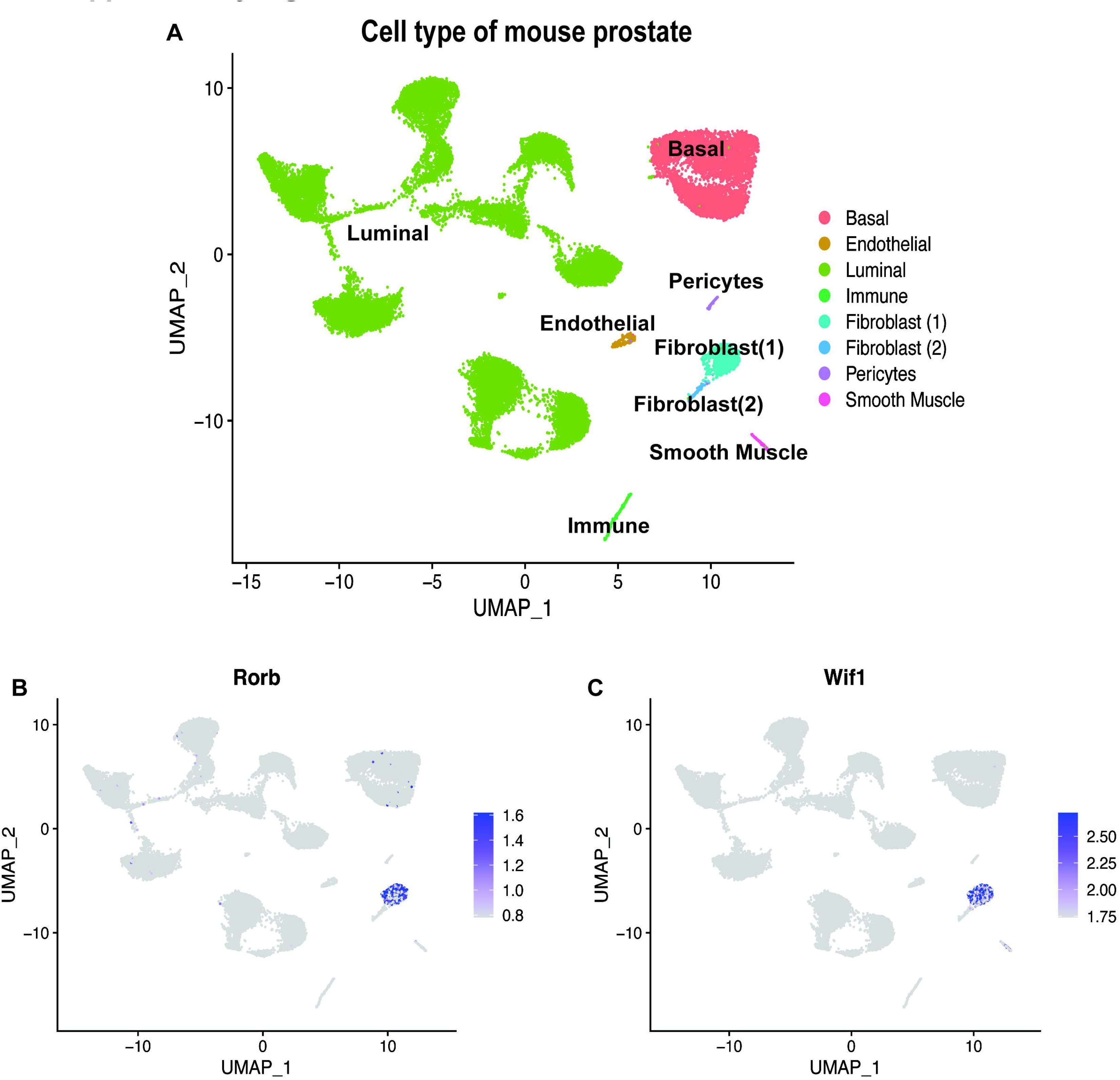
*Wif1* is expressed in a subset of mouse fibroblast. A) UMAPs of mouse prostate cell clusters and expression of B) *Rorb*, and C) *Wif1*.

**Supplementary Fig. 5.**
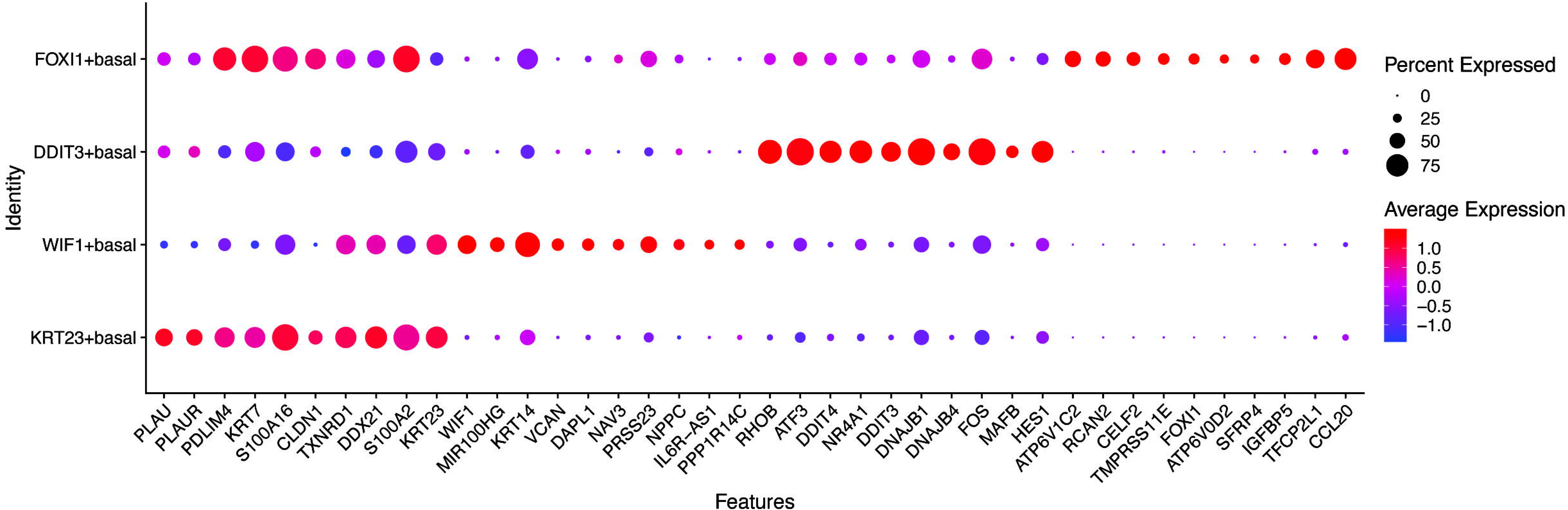
Dotplot of top 10 differentially expressed genes by adjusted p-value and log2FC for different basal cell subtypes of BPH and organ donor samples.

**Supplementary Fig. 6.**
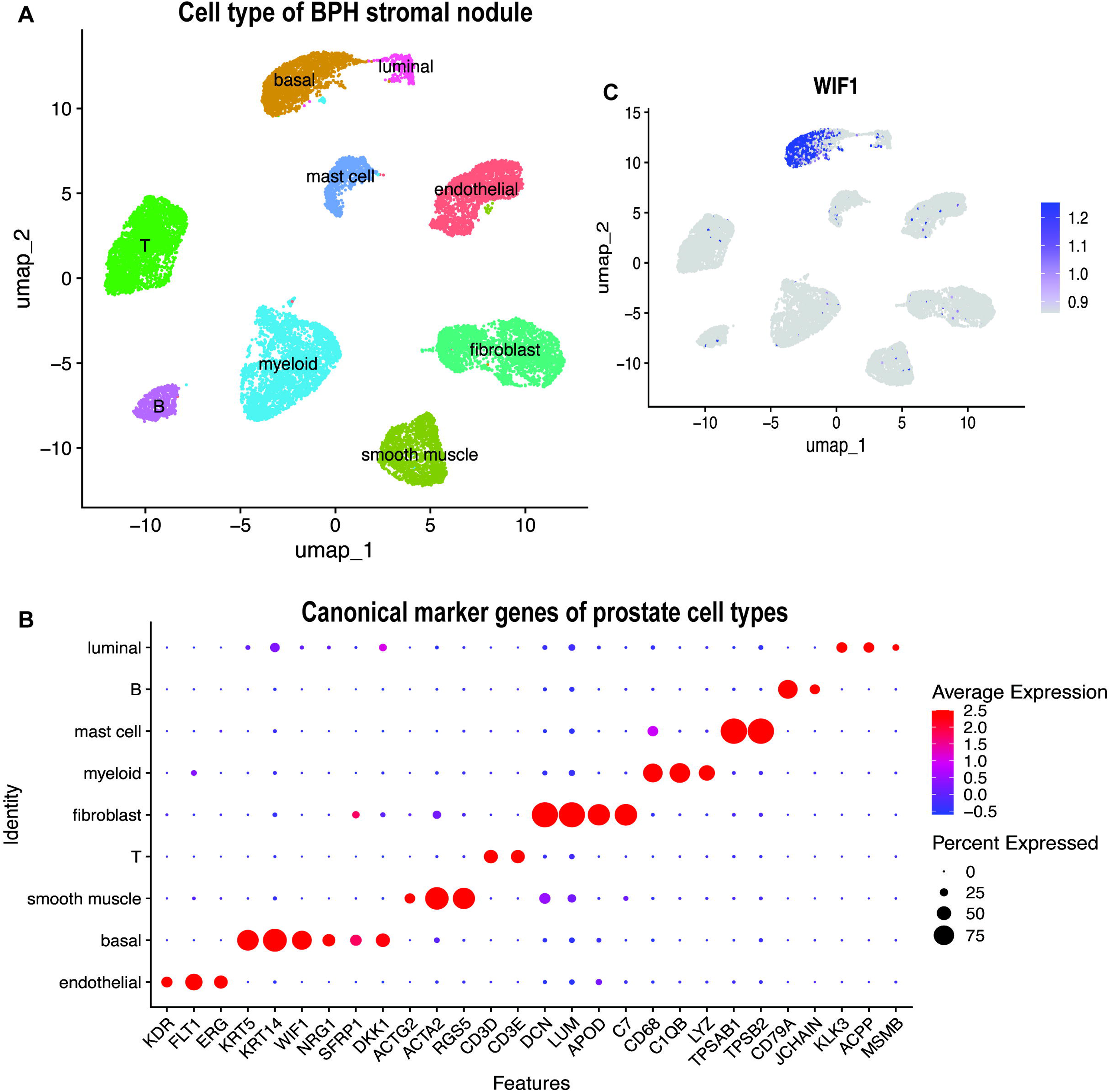
*WIF1*+ basal cell subtype is the major basal cell type in BPH stromal nodule sample. A) UMAP and clustering analysis of previously published scRNA-seq dataset of BPH stromal nodule samples by Joseph et al. [4], showed cell clustering by known cell types in the BPH stromal samples. B) Dotplot of expression of canonical marker genes of prostate cell types in BPH stromal samples. C) Expression of *WIF1* in BPH stromal samples.

**Supplementary Fig. 7.**
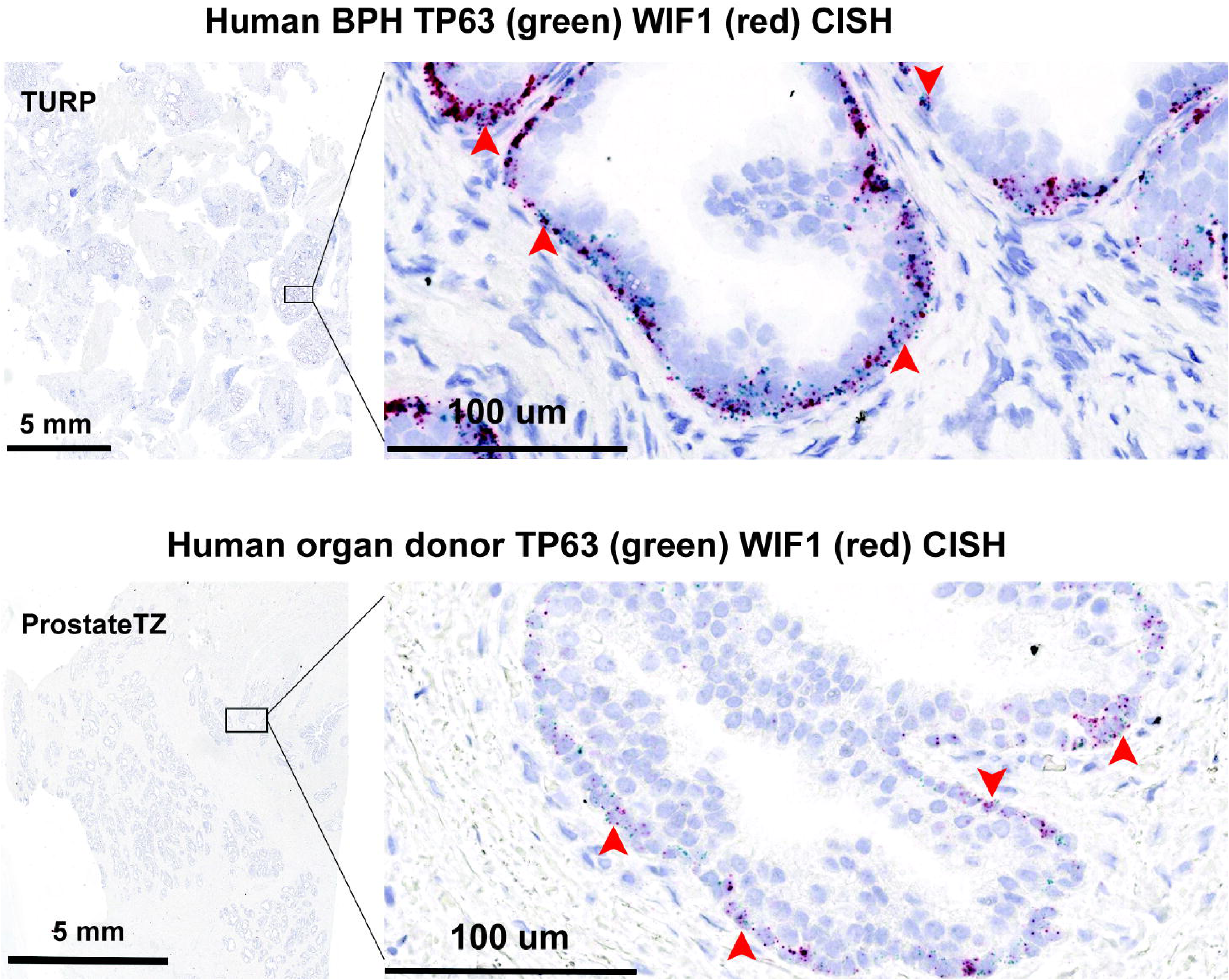
*WIF1*+ basal Cell is Enriched in BPH. Representative examples of CISH staining of *TP63* (green), a known basal cell marker, and *WIF1* (red), a *WIF1*+ basal cell subtype marker, in BPH and donor TZ samples. For 0.3x magnification, the scale bar indicates 5 mm. For 40x magnification, scale bar indicates 100 μm. The red arrow indicates dual expression of *WIF1* and *TP63* in the BPH and donor TZ samples.

**Supplementary Fig. 8.**
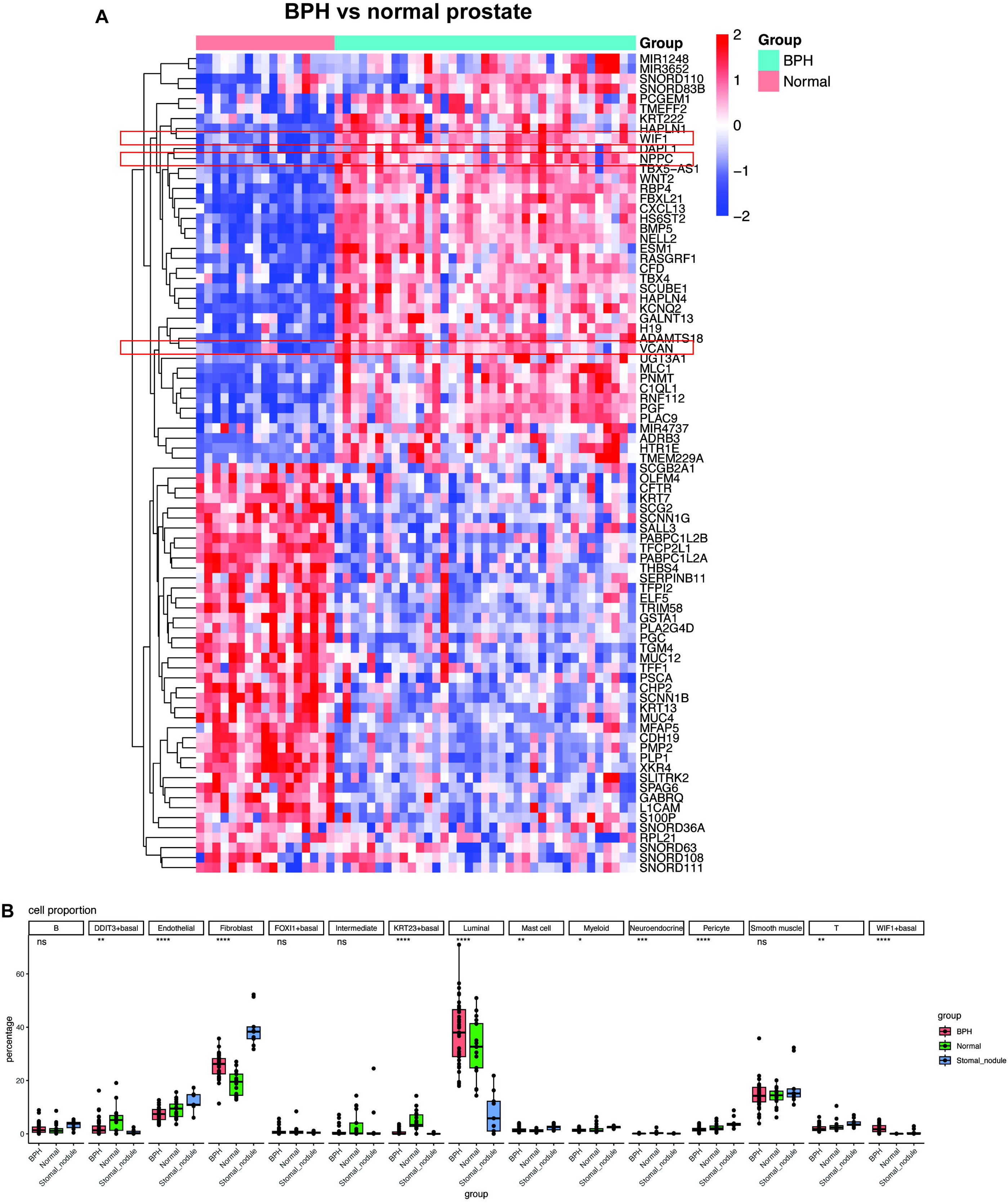
Enrichment of *WIF1*+ basal cells in BPH samples relative to both normal prostate and BPH stromal nodule of previously published bulk RNA-seq data by Middleton et al. [22]. A) Heatmap showing the most significant DEGs between BPH and normal prostate. Genes were selected by fold change > 5 with adjusted p-value < 0.05. *WIF1*+ basal specific genes-*WIF1, NPPC,* and *VCAN* were highlighted. B) Proportions of each cell type defined by our single-cell RNA-seq dataset were estimated by RCTD for each sample. Each dot represents one bulk RNA-seq sample. Box-whiskers are colored by group (BPH, Normal, BPH stromal nodule). Horizontal bars indicate mean cell proportions per group. P values were calculated using one-way ANOVA. ns: P > 0.05; *: P < 0.05; **: P < 0.01; ***: P < 0.001; ****: P < 0.0001.

